# Reference nodule transcriptomes for *Melilotus officinalis* and *Medicago sativa* cv. Algonquin

**DOI:** 10.1101/2022.02.08.479627

**Authors:** Rui Huang, Wayne A Snedden, George C diCenzo

## Abstract

Host/symbiont compatibility is a hallmark of the symbiotic nitrogen-fixing interaction between rhizobia and legumes, mediated in part by plant produced nodule-specific cysteine-rich (NCR) peptides and the bacterial BacA membrane protein that can act as a NCR peptide transporter. In addition, the genetic and metabolic properties supporting symbiotic nitrogen fixation often differ between compatible partners, including those sharing a common partner, highlighting the need for multiple study systems. Here, we report high quality nodule transcriptome assemblies for *Medicago sativa* cv. Algonquin and *Melilotus officinalis*, two legumes able to form compatible symbioses with *Sinorhizobium meliloti*. The compressed *M. sativa* and *M. officinalis* assemblies consisted of 79,978 and 64,593 contigs, respectively, of which 33,341 and 28,278 were assigned putative annotations, respectively. As expected, the two transcriptomes showed broad similarity at a global level. We were particularly interested in the NCR peptide profiles of these plants, as these peptides drive bacterial differentiation during the symbiosis. A total of 412 and 308 NCR peptides were predicted from the *M. sativa* and *M. officinalis* transcriptomes, respectively, with approximately 9% of the transcriptome of both species consisting of *NCR* transcripts. Notably, transcripts encoding highly-cationic NCR peptides (isoelectric point > 9.5), which are known to have antimicrobial properties, were ~2-fold more abundant in *M. sativa* than in *M. officinalis*, and ~27-fold more abundant when considering only NCR peptides in the six-cysteine class. We hypothesize that the difference in abundance of highly-cationic NCR peptides explains our previous observation that some rhizobial *bacA* alleles which can support symbiosis with *M. officinalis* are unable to support symbiosis with *M. sativa*.

## INTRODUCTION

Leguminous plants are able to establish symbiotic relationships with a group of soil bacteria known as rhizobia. During the interaction, the rhizobia are located within a specialized organ known as a nodule where they fix atmospheric nitrogen into ammonia in exchange for reduced carbon from their host. Symbiosis is initiated following an exchange of chemical signals in the rhizosphere between compatible partners (1): legumes secrete flavonoids that attract soil rhizobia and induce expression of rhizobial *nod* genes, leading to rhizobial production of chito-oligosaccharide Nod factors that elicit the nodulation process by legumes. This process involves the curling of root hairs to trap rhizobia, and the formation of infection threads within which rhizobia divide and move toward the root cortical layer (2). Rhizobia released from infection threads are endocytosed into the cytoplasm of nodule cells, where they develop into mature N_2_-fixing bacteroids. In some legumes, such as those belonging to the Inverted Repeat Lacking Clade (IRLC), the rhizobia undergo an irreversible host-induced process known as terminal differentiation that is largely driven by a unique class of legume proteins known as nodule-specific cysteine-rich (NCR) peptides (3). Terminal differentiation involves cell enlargement, genome endoreplication, and increased membrane permeability, and is thought to increase the efficiency of N_2_-fixation (4–6).

Not all rhizobium/legume pairings are compatible (7, 8). Partner compatibility is determined by numerous factors impacting both early and late stages of the symbiotic interaction (9). The flavonoids secreted by legumes vary, as does the ability of rhizobia to respond to different flavonoids (10–13). Similarly, the Nod factor produced by rhizobia differ and legume hosts respond only to Nod factors with specific structures (14). Moreover, legume infection depends on rhizobia producing particular host-compatible exopolysaccharide molecules (15, 16), and variations in exopolysaccharide structure can impact specificity at the level of plant ecotype and bacterial strain (17). In addition, some rhizobia secrete effector proteins that induce effector-triggered immune responses in a cultivar-specific manner, thereby influencing host range (18–20). Moreover, for IRLC legumes, an effective symbiotic interaction requires compatibility between the host-produced NCR peptides and the rhizobial membrane protein BacA (21–24).

NCR peptides are a large class of legume-specific proteins, with ~ 600 members in *Medicago truncatula* (25). These proteins display little conservation in amino acid composition but possess four or six cysteine residues at conserved positions (26). The length of mature NCR peptides varies from about 20 to 50 amino acids and includes two or three disulfide bridges (27). NCR peptides can be classified as either cationic (isoelectric point [pI] ≥ 8), neutral (6 ≤ pI < 8), or anionic (pI < 6) (27). Highly cationic peptides (pI ≥ 9.0) display antimicrobial activity *in vitro*, likely through disrupting microbial membranes, thereby leading to permeabilization and cell lysis (28, 29). *In planta*, NCR peptides are required for rhizobium terminal differentiation and an effective symbiosis in IRLC legumes (3). Deletion of individual *NCR* genes is sufficient to block N_2_-fixation (30, 31); however, mutation of other *NCR* genes can result in N_2_-fixation in previously incompatible symbioses (21, 22), demonstrating the role of NCR peptides in partner compatibility.

The ability of rhizobia to establish an effective symbiosis with IRLC legumes requires the membrane protein BacA (32). BacA functions as a peptide transporter (33), and *bacA* deletion mutants are both unable to import NCR peptides and show increased sensitivity to cationic NCR peptides (34–36). In addition, rhizobium *bacA* mutants are unable to fix nitrogen in symbiosis with IRLC legumes; instead, the rhizobia are quickly killed in a NCR peptide-dependent fashion upon release from the infection threads (32, 34). Intriguingly, BacA appears to be a host-range determinant factor in IRLC legumes. For example, studies have shown that introduction of the *bacA* or *bacA*-like genes of *Mesorhizobium loti* and *Bradyrhizobium* species into a *Sinorhizobium meliloti bacA* mutant is insufficient to allow N_2_-fixation during interaction with IRLC legumes of the genus *Medicago* (35, 37). Similarly, we previously demonstrated that replacement of the *S. meliloti bacA* with the *bacA* alleles of *Sinorhizobium fredii* NGR234 or *Rhizobium leguminosarum* bv. *viciae* 3841 does not allow for N_2_-fixation during symbiosis with *Medicago sativa* (alfalfa) but does support N_2_-fixation on the IRLC legumes *Melilotus alba* (white sweet clover) and *Melilotus officinalis* (yellow sweet clover) ((24) and **Table S1**).

In addition to the above-noted comparison, several symbiotic differences have been observed when *S. meliloti* mutants interact with *Medicago* versus *Melilotus* plants (38–41), suggesting that *Melilotus* plants are a valuable secondary model system to study the symbiotic properties of *S. meliloti*. To further develop *M. officinalis* as a model species for studying symbiosis, here we report a reference nodule transcriptome for *M. officinalis*. We further compare the characteristics and the expression of *NCR* genes between *M. officinalis* and *M. sativa* to investigate whether the ability of certain *bacA* alleles to support symbiosis with *Melilotus* but not *Medicago* plants is correlated with differences in the NCR peptide profile of these genera.

## MATERIALS AND METHODS

### Plant materials and sample collection

*M. sativa* cv. Algonquin (alfalfa) and *M. officinalis* (yellow blossom sweet clover) seeds were purchased from Speare Seeds Limited (Harriston, Ontario, Canada). Seeds were surface sterilized with 95% ethanol for five minutes followed by 2.5% hypochlorite for 20 minutes, and then soaked in sterile double-distilled water (ddH_2_O) for one hour. The sterilized seeds were plated on 1X water agar plates and incubated at room temperature in the dark for two days. Five germinated seeds were placed in autoclaved Leonard Assemblies consisting of two Magenta Jars with a cotton wick extending from the top jar (containing vermiculiate mixed with silica sand [1:1 *w/w*]) into the bottom jar (containing 250 mL of Jensen’s media (42)), and then incubated in a Conviron growth chamber for two nights. Wildtype *S. meliloti* strain Rm2011 was grown overnight at 30°C in LBmc broth (10 g L^-1^ tryptone, 5 g L^-1^ yeast extract, 5 g L^-1^ NaCl, 2.5 mM CaCl_2_, and 2.5 mM MgCl_2_), washed with 0.85% NaCl, and diluted to a density of ~ 1×10^7^ CFU mL^-1^ in sterile ddH_2_O. Ten mL of cell suspension was then added to each Leonard Assembly. Plants were grown in a Conviron growth chamber with a day (18 hours, 21°C, light intensity of 300 µmol m^-2^ s^-1^) and night (6 hours, 17°C) cycle. Root nodules were collected four weeks post-inoculation and immediately flash frozen with liquid N_2_ and stored at –80°C until use. All nodules collected from plants grown in the same Leonard Assembly were stored in a single tube and treated as one replicate. The shoots from each pot were dried at 60°C for two weeks prior to measuring shoot dry weight (**Table S2**).

### RNA extraction and sequencing

Total RNA from three replicates of frozen *M. sativa* and *M. officinalis* nodule tissue was extracted using Direct-zol RNA miniprep kits (ZYMO Research) according to the manufacturer’s protocol. Total RNA samples were treated with DNase I (New England Biolabs) to degrade any contaminating DNA according to the manufacturer’s protocol, and the RNA again purified using Direct-zol RNA miniprep kits. Total RNA samples were run on a MOPS-formaldehyde agarose gel (119 mL MOPS buffer [200 mM MOPS, 80 mM sodium acetate, 10 mM EDTA, pH 7.0, in DEPC-treated ddH_2_O], 6 mL formaldehyde, 1.25 g agarose) to check the integrity of the RNA (**Figure S1**), and subsequently verified using an Agilent Bioanalyzer chip.

Library preparation and Illumina sequencing were performed at The Centre for Applied Genomics at The Hospital for Sick Children (Toronto, Ontairo, Canada). Libraries were prepared using the NEB Next^®^ Ultra™ II Directional RNA Library Prep Kit for Illumina^®^. Libraries were then sequenced using one lane of a high throughput flow cell on an Illumina HiSeq 2500 platform, generating 125 bp paired-end reads.

### Transcriptome *de novo* assembly and quality control

The nodule transcriptomes of *M. sativa* and *M. officinalis* were *de novo* assembled following the same procedure. First, reads from the triplicate samples were combined, and then preprocessing of the raw reads was performed to ensure only high-quality data was used for *de novo* transcriptome assembly. Read quality was initially evaluated using FastQC version 0.11.9 (43), following which errors in raw reads were identified and corrected by the k-mer based method of Rcorrector version 1.0.4 (44). The FilterUncorrectablePEfastq.py script (github.com/harvardinformatics/TranscriptomeAssemblyTools/) was used to remove any read pair where at least one read had an unfixable error identified by Rcorrector. Adaptors sequences, short reads (< 25 bp), and low-quality reads (Q score < 20) were removed using Trim Galore version 0.6.6 (bioinformatics.babraham.ac.uk/projects/trim_galore/), which is a wrapper calling cutadapt version 3.2 (45) and FastQC (**Table 1**). The processed reads were further trimmed by Trimmomatic version 0.4.0 (46) included in the Trinity software distribution with the following parameters: *SLIDINGWINDOW:5:20 LEADING:3 TRAILING:3 MINLEN:25*. The quality and presence of adaptors in the preprocessed reads were then examined using FastQC. Following preprocessing, 174,707,055 and 119,333,821 paired end reads (~43.7 and ~29.8 Gb, respectively) remained for *M. sativa* and *M. officinalis*, respectively (**Table S3**).

**Table 1.**
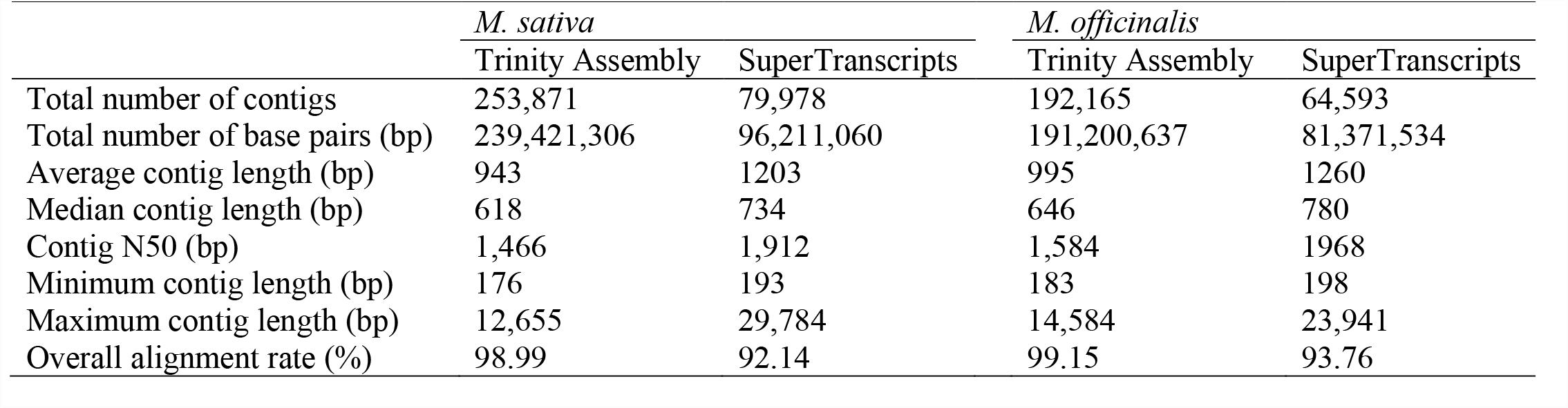
Summary statistics from the *de novo* Trinity and compressed (SuperTranscripts) nodule transcriptome assemblies.

The preprocessed reads were assembled using Trinity version 2.9.0 without genome guidance (47). Then, the assembled contigs were clustered into gene-level clusters using SuperTranscripts (48). Gene isoforms were identified by Corset version 1.09 with the log likelihood ratio threshold set to very high (49). Based on the Corset clusters, Lace version 1.14.1 was used to merge the gene isoforms into single long supertranscripts meant to provide a gene-like view of the transcriptome (48).

Multiple methods were used to examine the quality of the Trinity and SuperTranscript assemblies. First, the alignment rates of the preprocessed reads to the assemblies were inspected using STAR version 2.7.8a with the two-pass mode that is more sensitive to alternative splicing (50). Second, assembly statistics such as N50 and number of contigs were calculated using the seqstats software (github.com/clwgg/seqstats). Third, the completeness of the assembies was evaluated using BUSCO version 5.1.2, run separately using the OrthoDB v10 ‘Fabales’ and ‘Viridiplantae’ reference databases (51). The assemblies were also compared to the *S. meliloti* Rm2011 genome (52) using BLASTn version 2.5.0+ (53), which confirmed the absence of contaminating *S. meliloti* transcripts in the assemblies. Finally, the *M. sativa de novo* assembly was aligned to a publicly-available genome of *M. sativa* cultivar XinJiangDaYe (54) with MUMmer version 4.0+, and 87.3% of transcripts were sucessfully aligned to the genome.

### Transcriptome annotation

Coding regions within the supertranscripts were predicted by TransDecoder version 5.5.0 (<u>github.com/TransDecoder/TransDecoder</u>), using the results of BLASTp searches (E-value cutoff of 1e-5) against the Uniport database as ORF retention criteria (2021 January release) (55). The functional annotation of the predicted coding sequences then proceeded via three steps. First, BLAST bidirectional best hits between the *M. truncatula* A17 proteome (assembly release r5.0 1.7) (56) and the longest predicted protein isoform of each contig of our transcriptome assemblies were identified using BLASTp (E-value cutoff of 1e-5, culling limit 1). For all bidirectional best hits, the annotations from *M. truncatula* A17 were transferred to the corresponding contigs of the *M. sativa* or *M. officinalis* transcriptome. Second, all predicted protein isoforms of each contig in each transcriptome assembly were annotated using eggNOG-mapper version 2.1.0 with DIAMOND version 2.0.4 and the Viridiplantae dataset (E-value cutoff of 1e-3) (57, 58). Third, for each conting not annotated by BLAST or eggNOG-mapper, the hmmsearch function of HMMER version 3.3.2 was used to search all predicted protein isoforms against the complete set of hidden Markov models (HMMs) from the Pfam version 34.0 database and separately against the TIGRFAM version 15.0 HMM database (E-value cutoff of 1e-5) (59–61), and results were filtered to remove annotations with a Bit-score < 50. For repetitive annotations from isoforms of a gene, only the consensus annotations were retained. For contigs successfully annotated by more than one of the annotation methods, results from the bidirectional BLAST took priority, followed by the results of eggNOG-mapper, then the Pfam searches, and finally the TIGRFAM searches.

### NCR peptide identification

Considering the high degree of sequence diversity of NCR peptide sequences, the functional annotation methods described above were not sufficiently sensitive to discover genes encoding NCR peptides in the assemblies. Therefore, the SPADA version 1.0 pipeline was used to identify NCR peptides (62). SPADA is specialized to predict cysteine-rich peptides in plant genomes and is distributed with a *M. truncatula* prediction model. Cysteine-rich peptides in the *M. sativa* and *M. officinalis* assemblies were predicted using the SPADA pipeline with following software: HMMER version 3.0, Augustus version 2.6, GeneWise version 2.2.0, GeneMark.hmm eukaryotic version 3.54, GlimmerHMM version 3.0.1, and GeneID version 1.1 (63–66). The putative NCR peptide sequences were filtered to remove those without a signal peptide, and then further filtered based on the E-value (cutoff of 1e-5) and hmm score (cutoff of 50). Filtered sequences were then verified via hmmscan searches against the Pfam database, and they were aligned using Clustal Omega version 1.2.4 (67) to ensure the presence of the signature cysteine motif and N terminal signal peptide that are present in *bona fide* NCR peptides.

### NCR peptides classification

To predict the lengths of mature NCR peptides, signalP version 4.1g with the notm network was used to predicted cleavage sites and extract mature NCR peptides (68), and the number of cysteine residues in each motif were counted. The pI values of the NCR peptides were predicted using the pIR R package, and the value for each peptide was calculated based on the mean values from all prediction methods excluding the highest and lowest values (69).

### Gene-expression level estimation and differential expression analysis

Gene-expression levels of each *M. sativa* and *M. officinalis* replicate transcriptome were estimated by transcript abundance estimation using salmon version 0.12.0 in mapping-based mode (library type automatic, validate Mapping) (70) and the reference transcriptomes produced as described above. R package deseq2 version 1.32.0 (71) was used to perform differential expression analysis between *M. sativa* and *M. officinalis*, using the raw counts from salmon, the length of each gene in each species as an additional parameter during normalization, and limiting the analysis to one-to-one orthologs identified by OrthoFinder version 2.5.2 (72). OrthoFinder was run with default settings using the total predicted *M. sativa* and *M. officinalis* proteomes including all isoforms, following which orthologs were reduced to one per supertransript.

### Gene Ontology term analysis

The Gene Ontology (GO) terms for *M. sativa* and *M. officinalis* were obtained from the *M. truncatula* A17 proteome (assembly release r5.0 1.7) and annotations from eggNOG-mapper. For transcripts annotated with GO terms from both sources, the concensus GO term annotations were retained. Then, the GO terms were reduced based on the Generic GO subset (download 10 August 2021).

### Software information

All analyses were performed in an Ubuntu 20.04.2 LTS (Linux 5.8.0-48-generic) operation system or on the Compute Canada Graham cluster. Custom scripts were written in Python version 3.8.5 and bash. R version 3.6.3 was used during data analysis (73).

### Data availability

All custom scripts to perform the analyses described in this study are available through GitHub (https://github.com/hyhy8181994/Nodule_transcriptome_script). Raw Illumina data are available through the Short Read Archive (SRR15724671, SRR15724670, SRR15724669, SRR15724668, SRR15724667, and SRR15724666) hosted by the National Center for Biotechnology Information (NCBI). The assembled transcriptomes are available through the Transcriptome Shotgun Assembly Sequence Database (GJLW00000000 and GJLK00000000) hosted by the NCBI.

## RESULTS AND DISCUSSION

### Reference nodule transcriptomes for *Melilotus officinalis* and *Medicago sativa* cv. Algonquin

To establish reference nodule transcriptomes of *M. sativa* cv. Algonquin and *M. officinalis* during symbiosis with *S. meliloti* Rm2011, the poly-A enriched RNA from triplicate samples was sequenced using Illumina technology (2×125 bp paired-end reads), generating ~50 Gb (~ 202 million paired-end reads) and ~35 Gb (~139 million paired-end reads) of data for *M. sativa* and *M. officinalis*, respectively (see **Table S3** for sequencing statistics). *De novo* assembly of the *M. sativa* sequencing data resulted in 253,871 contigs, while 192,165 *de novo* assembled contigs were produced for *M. officinalis*. Contigs expected to represent splice variants of a single gene were merged into so-called “supertranscripts” using the SuperTranscripts program, resulting in compressed assemblies of 79,978 and 64,593 contigs for *M. sativa* and *M. officinalis*, respectively (**Table 1**). Transcriptomes were annotated as described in the Materials and Methods, resulting in putative annotations for 33,431 *M. sativa* contigs and 28,278 *M. officinalis* contigs (**Datasets S1 and S2**). Of these, ~ 52% (*M. sativa*) and ~ 58% (*M. officinalis*) are high confidence annotations as they were transferred from the *M. truncatula* whole genome annotation following identification of putative orthologs using a BLAST bidirectional best hit approach (**Table S4**). Considering that previous studies have predicted the presence of ~23,000 long non-coding RNAs (lncRNAs) in *M. truncatula* (74) and ~47,000 lncRNAs in the legume *Pisum sativum* (pea) (75), we hypothesize that the majority of the unannotated *M. sativa* and *M. officinalis* transcripts reflect lncRNAs.

All of the examined assembly summary statistics (mean and median contig length, contig N50) were improved in the compressed assemblies compared to the original *de novo* assemblies, indicating that the compressed assemblies are of higher structural quality (**Table 1**). The *M. sativa* transcriptome summary statistics, such as N50 and and transcript length, are consistent with those reported for other *M. sativa de novo* transcriptome assemblies, although the number of transcripts varies likely due to each study examining different tissues (76, 77). In addition, the assemblies appear to be robust; greater than 90% of the filtered reads used for transcriptome assembly could be mapped to the corresponding assemblies by STAR (**Table 1**). Moreover, > 92% and > 83% of the Viridiplantae and Fabales BUSCO marker genes, respectively, were identified as complete and single-copy in the *M. sativa* and *M. officinalis* compressed assemblies (**Figure 1**). The structural quality (e.g. high average and median contig length and N50) and BUSCO benchmark scores described here are in line with those reported for other plant *de novo* transcriptome assemblies (78–80). Taken together, these results indicate that our *M. sativa* cv. Algonquin and *M. officinalis* reference nodule transcriptomes are reliable and of high quality.

**Figure 1.**
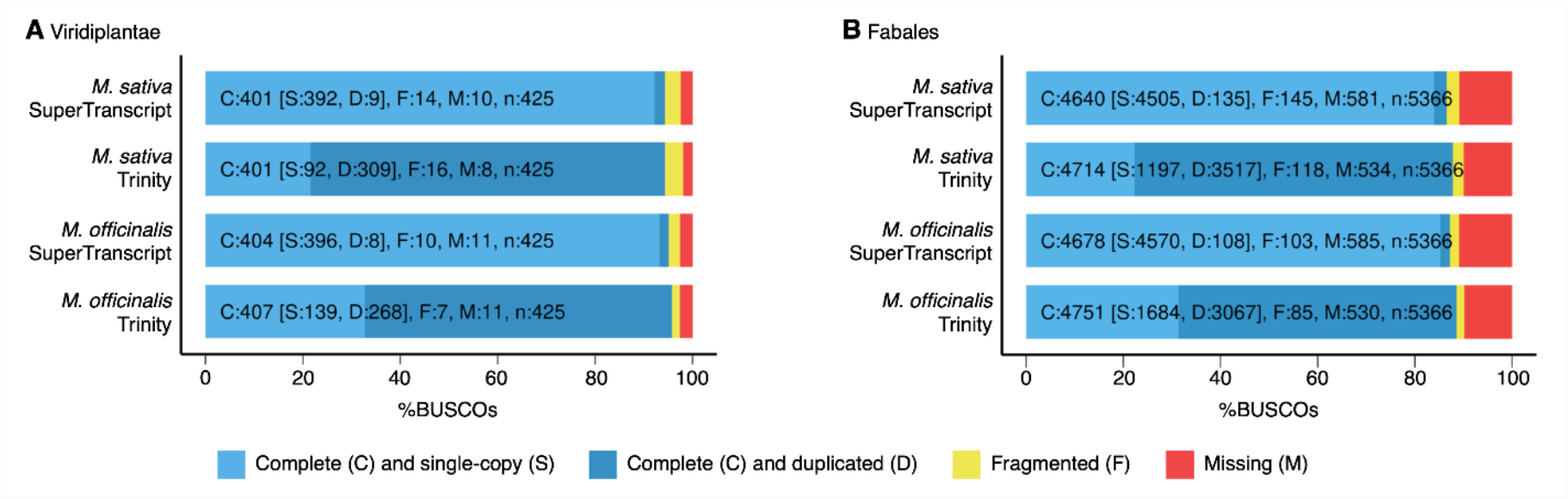
Estimates of nodule transcriptome completeness. Completeness of the *M. sativa* and *M. officinalis* nodule transcriptome assemblies was assessed using BUSCO with the (**A**) Viridiplantae and (**B**) Fabales single-copy marker gene datasets. The fraction of BUSCO genes identified as complete and single-copy (light blue), complete but duplicated (dark blue), fragmented (yellow), and missing (red) is shown.

### Comparative transcriptome analysis between *M. sativa* and *M. officinalis*

As an initial examination of the *M. sativa* and *M. officinalis* transcriptomes, the annotated functions of the proteins predicted to be encoded by the supertranscripts were summarized using the Generic GO term subset (**Figure 2, Dataset S3 and S4**). Approximately 18,363 (26.1%) of the *M. sativ*a supertranscripts and 16,674 (28.6%) of the *M. officinalis* supertranscripts were annotated with GO terms. No significant difference in the GO term profiles of the two species was observed, with the five most frequently annotated biological process GO terms being GO:0008150 (biological process), GO:0006950 (response to stress), GO:0006464 (cellular protein modification process), GO:003464 (cellular nitrogen compound metabolic process) and GO:0048856 (anatomical structure development). At this broad scale, the GO term data suggest that there is substantial similarity in the nodule transcriptomes of *M. sativ*a and *M. officinalis*.

**Figure 2.**
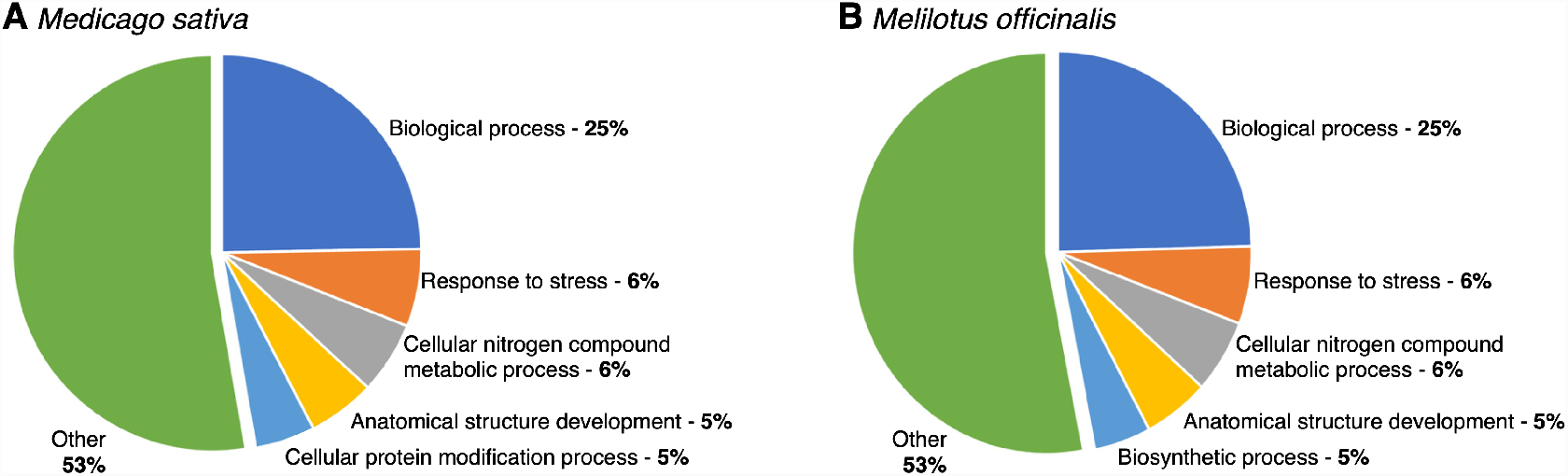
Summary of the Slim GO Biological Processes annotations for the nodule transcriptomes. Transcripts were annotated with Slim GO terms, and the annotations for the biological processes were summarized as pie charts for (**A**) *M. sativa* and (**B**) *M. officinalis*.

We next examined the predicted functions of the proteins encoded by the 50 most abundant transcripts in both species (**Table 2 and 3**). Not surprisingly, these transcripts were enriched in those predicted to encode nodulins and leghaemoglobin-like proteins. Nodulins refer to diverse proteins expressed specifically in nodule tissue, which play various structural or metabolic roles during symbiotic nitrogen fixation. Among the nodulins, are the leghemoglobin proteins that account for up to 40% of the total soluble protein in legume nodules (81). Leghemoglobins play an important role in maintaining the low free-oxygen concentration required to protect the oxygen-sensitive nitrogenase enzyme (82).

**Table 2.** The 50 most highly abundant transcripts in the *M. sativa* nodule transcriptome, with the average expression level in transcripts per million (TPM) and the functional annotation.

| Gene ID | TPM | Functional Prediction |
| --- | --- | --- |
| Cluster-2.43518 | 30,662 | hypothetical protein (hypothetical leghaemoglobin) |
| Cluster-2.26078 | 17,769 | Putative albumin I |
| Cluster-2.23447 | 12,515 | hypothetical protein (hypothetical leghaemoglobin) |
| Cluster-2.23176 | 8,065 | Nodulin-25 |
| Cluster-2.22272 | 7,992 | Putative Late nodulin |
| Cluster-2.23033 | 6,778 | None |
| Cluster-2.21873 | 5,919 | hypothetical protein |
| Cluster-2.22935 | 5,253 | Belongs to the globin family |
| Cluster-2.33070 | 5,232 | Putative ribonuclease H-like domain-containing protein |
| Cluster-2.24197 | 4,245 | Component of the replication protein A complex (RPA) |
| Cluster-2.22983 | 4,196 | Belongs to the globin family |
| Cluster-2.24546 | 3,886 | Predicted NCR peptide (crp1450_Cluster-2.24546_0M_1) |
| Cluster-2.19511 | 3,344 | None |
| Cluster-2.26207 | 3,296 | Extensin-like_protein_repeat |
| Cluster-2.49512 | 3,173 | Putative Blue (type 1) copper binding protein |
| Cluster-2.21810 | 3,050 | Predicted NCR peptide (crp1160_Cluster-2.21810_0M_1) |
| Cluster-2.22936 | 3,014 | Belongs to the globin family |
| Cluster-2.29430 | 2,789 | Putative Late nodulin |
| Cluster-2.22881 | 2,788 | Predicted NCR peptide (crp1430_Cluster-2.22881_0M_1) |
| Cluster-2.22836 | 2,509 | None |
| Cluster-2.23729 | 2,271 | hypothetical protein |
| Cluster-2.22458 | 2,245 | Belongs to the globin family |
| Cluster-2.24829 | 2,227 | hypothetical protein |
| Cluster-2.25168 | 2,209 | None |
| Cluster-2.24278 | 2,093 | Nodule-specific_GRP_repeat |
| Cluster-2.23928 | 2,038 | Late_nodulin_protein |
| Cluster-2.22042 | 2,029 | None |
| Cluster-2.23245 | 2,019 | Putative translationally controlled tumor protein |
| Cluster-2.25794 | 2,018 | Putative protein-synthesizing GTPase |
| Cluster-2.31376 | 1,957 | Predicted NCR peptide (crp1190_Cluster-2.31376_0M_1) |
| Cluster-2.21809 | 1,939 | Predicted NCR peptide (crp1160_Cluster-2.21809_0M_1) |
| Cluster-2.28083 | 1,854 | Predicted NCR peptide (crp1210_Cluster-2.28083_0M_1) |
| Cluster-2.23524 | 1,844 | Early nodulin-16 |
| Cluster-2.34649 | 1,839 | Putative Late nodulin |
| Cluster-2.16993 | 1,808 | hypothetical protein |
| Cluster-2.21813 | 1,731 | None |
| Cluster-2.22457 | 1,705 | Belongs to the globin family |
| Cluster-2.28283 | 1,674 | None |
| Cluster-2.26205 | 1,621 | Predicted NCR peptide (crp1240_Cluster-2.26205_0M_1) |
| Cluster-2.23310 | 1,607 | asparagine synthetase |
| Cluster-2.21870 | 1,596 | Nodule-specific_GRP_repeat |
| Cluster-2.19604 | 1,583 | Predicted NCR peptide (crp1420_Cluster-2.19604_0M_1) |
| Cluster-2.18485 | 1,509 | hypothetical protein |
| Cluster-2.26876 | 1,458 | Predicted NCR peptide (crp1410_Cluster-2.26876_0M_1) |
| Cluster-2.21536 | 1,441 | Ubiquitin_family |
| Cluster-2.22785 | 1,408 | None |
| Cluster-2.23374 | 1,385 | Predicted NCR peptide (crp1420_Cluster-2.23374_0M_1) |
| Cluster-2.28787 | 1,311 | Putative Late nodulin |
| Cluster-2.22402 | 1,292 | Predicted NCR peptide (crp1520_Cluster-2.22402_0M_1) |
| Cluster-2.30081 | 1,289 | Late nodulin protein |

**Table 3.** The 50 most highly abundant transcripts in the *M. officinalis* nodule transcriptome, with the average expression level in transcripts per million (TPM) and the functional annotation.

| Gene ID | TPM | Functional Prediction |
| --- | --- | --- |
| Cluster-3554.18801 | 38,953 | Belongs to the globin family |
| Cluster-3554.18778 | 14,091 | Belongs to the globin family |
| Cluster-3554.16063 | 10,412 | Late_nodulin_protein |
| Cluster-3554.15387 | 7,885 | Putative Late nodulin |
| Cluster-3554.16088 | 6,146 | Putative Late nodulin |
| Cluster-3554.18892 | 6,104 | None |
| Cluster-3554.15771 | 5,813 | Late_nodulin_protein |
| Cluster-3554.18802 | 5,362 | Belongs to the globin family |
| Cluster-3554.18596 | 5,347 | Predicted NCR peptide (crp1430_Cluster-3554.18596_0M_1) |
| Cluster-3554.15456 | 3,912 | Putative translationally controlled tumor protein |
| Cluster-3554.18808 | 3,864 | Belongs to the globin family |
| Cluster-3554.33215 | 3,302 | hypothetical protein |
| Cluster-3554.18775 | 3,097 | Putative Late nodulin |
| Cluster-3554.23000 | 2,990 | Two predicted NCR peptide (crp1180_Cluster-3554.23000_0M_1 and crp1180_Cluster-3554.23000_0M_2) |
| Cluster-3554.36681 | 2,877 | Predicted NCR peptide (crp1500_Cluster-3554.36681_0M_1) |
| Cluster-3554.21555 | 2,868 | Predicted NCR peptide (crp1430_Cluster-3554.21555_0M_1) |
| Cluster-3554.16074 | 2,809 | Putative BURP domain-containing protein |
| Cluster-3554.29297 | 2,784 | hypothetical protein |
| Cluster-3554.18196 | 2,723 | None |
| Cluster-3554.18256 | 2,571 | Predicted NCR peptide (crp1440_Cluster-3554.18256_0M_1) |
| Cluster-3554.27063 | 2,522 | None |
| Cluster-3554.22577 | 2,449 | Putative Blue (type 1) copper binding protein |
| Cluster-3554.15361 | 2,409 | eEF1A |
| Cluster-3554.21311 | 2,384 | Belongs to the globin family |
| Cluster-3554.18838 | 2,340 | Late_nodulin_protein |
| Cluster-3554.18700 | 2,296 | Late_nodulin_protein |
| Cluster-3554.11713 | 2,261 | None |
| Cluster-3554.18706 | 2,259 | None |
| Cluster-3554.23445 | 2,214 | None |
| Cluster-3554.17366 | 2,141 | Predicted NCR peptide (crp1430_Cluster-3554.17366_0M_1) |
| Cluster-3554.23502 | 2,073 | hypothetical protein |
| Cluster-3554.25153 | 2,009 | Metallothionein-like protein 2 |
| Cluster-3554.21451 | 1,953 | Belongs to the globin family |
| Cluster-3554.28600 | 1,931 | hypothetical protein |
| Cluster-3554.22971 | 1,849 | hypothetical protein |
| Cluster-3554.18877 | 1,812 | Belongs to the glyceraldehyde-3-phosphate dehydrogenase family |
| Cluster-3554.13172 | 1,732 | Predicted NCR peptide (crp1440_Cluster-3554.13172_0M_1) |
| Cluster-3554.18195 | 1,704 | Zinc_knuckle |
| Cluster-3554.30751 | 1,653 | Putative Late nodulin |
| Cluster-3554.21774 | 1,635 | Late_nodulin_protein |
| Cluster-3554.13784 | 1,629 | Predicted NCR peptide (crp1420_Cluster-3554.13784_0M_1) |
| Cluster-3554.24033 | 1,611 | Prolyl isomerase (PPIase) |
| Cluster-3554.30294 | 1,556 | metallothionein-like protein |
| Cluster-3554.24764 | 1,552 | None |
| Cluster-3554.18368 | 1,552 | Nucleoside diphosphate kinase 1 |
| Cluster-3554.25344 | 1,537 | Metallothionein-like protein 1 |
| Cluster-3554.23646 | 1,514 | None |
| Cluster-3554.9874 | 1,512 | Belongs to the universal ribosomal protein uL13 family |
| Cluster-3554.18476 | 1,503 | Late_nodulin_protein |
| Cluster-3554.31722 | 1,476 | Putative Late nodulin |

To facilitate further comparison of the *M. sativa* and *M. officinalis* transcriptomes, the proteins predicted to be encoded by the supertranscripts of both species were arranged into orthologous groups using OrthoFinder. A total of 20,237 orthologous groups, accounting for 26,304 *M. sativa* and 24,895 *M. officinalis* supertranscripts, were identified. Interestingly, the abundance of the conserved supertranscripts was significantly higher, on average, than that of the species-specific transcripts (p value < 2.2e-16; **Figure 3**). In both plant species, the majority of the most abundant, species-specific annotated transcripts were also nodulins, globin family proteins that are likely species-specific leghaemoglobin isoforms, and some housekeeping genes such as ribonuclease and ribosomal proteins. It is noteworthy that the most abundant *M. sativa-*specific supertranscript is predicted to encode albumin I. Similarly, *M. officinalis* also has a highly-expressed albumin I supertranscript. The albumin I peptide family is known to be highly expressed in legume seeds and play roles in seed protection (83). Expression of albumin I genes has also been observed in *M. truncatula* root nodules, with expression specific to uninfected cells in the nitrogen fixation zone (84). These cells are thought to play essential roles in metabolite transport during symbiosis, and albumin I may have a role in protecting some of the nodule cells from rhizobium infection (84). A phylogenetic analysis of *M. truncaula* nodulins and albumin I peptides indicated that the *M. truncaula* albumin I clustered with a subset of nodulins, reflecting a close evolutionary relationship between these proteins (85).

**Figure 3.**
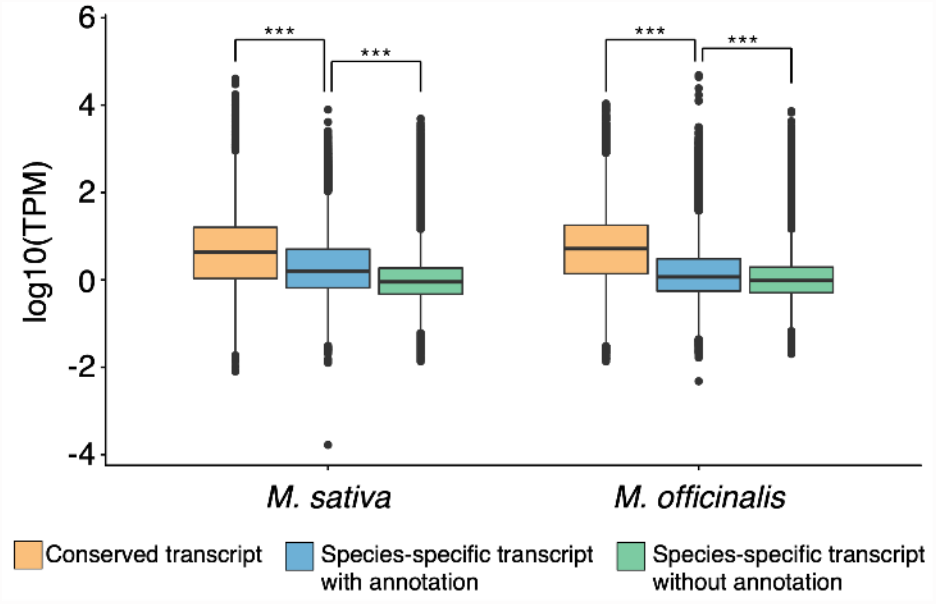
Transcript abundances for conserved and species-specific transcripts. Box plots displaying the distribution of average transcript abundances from triplicate samples, shown separately for genes with orthologs in both *M. sativa* and *M. officinalis* (orange), annotated transcripts found in only *M. sativa* or *M. officinalis* (blue), or transcripts that lack annotations and are found in only *M. sativa* or *M. officinalis* (green). Statistically significant differences between the distributions of a species are indicated with the asterisks (p-value < 1e^-10^; pairwise Wilcox tests).

We next compared the abundances of supertranscripts conserved in both *M. sativa* and *M. officinalis*, limiting the analysis to the 15,287 one-to-one orthologs detected by OrthoFinder. Desipite significant variation in the abundance of orthologous transcripts between *M. sativa* and *M. officinalis* – which may reflect limitations of inter-species transcriptome analysis – a clear correlation in the abundance of orthologous transcripts was detected (residual standard error = 0.517; **Figure 4**). Considering the limtations of inter-species differential expression analyses, we restricted our investigation to supertranscripts with absolute log_2_ fold changes > 5 and a p-value < 0.05. Using these thresholds, we identified 290 differentially-abundant transcripts, 86 of which were more abundant in *M. sativa*, and 204 of which were more abundant in *M. officinalis*. It should be noted, however, that only 159 of the differentially-abundant transcripts were annotated with the same or similar function in both species, and we focus on these 159 transcripts in the following discussion.

**Figure 4.**
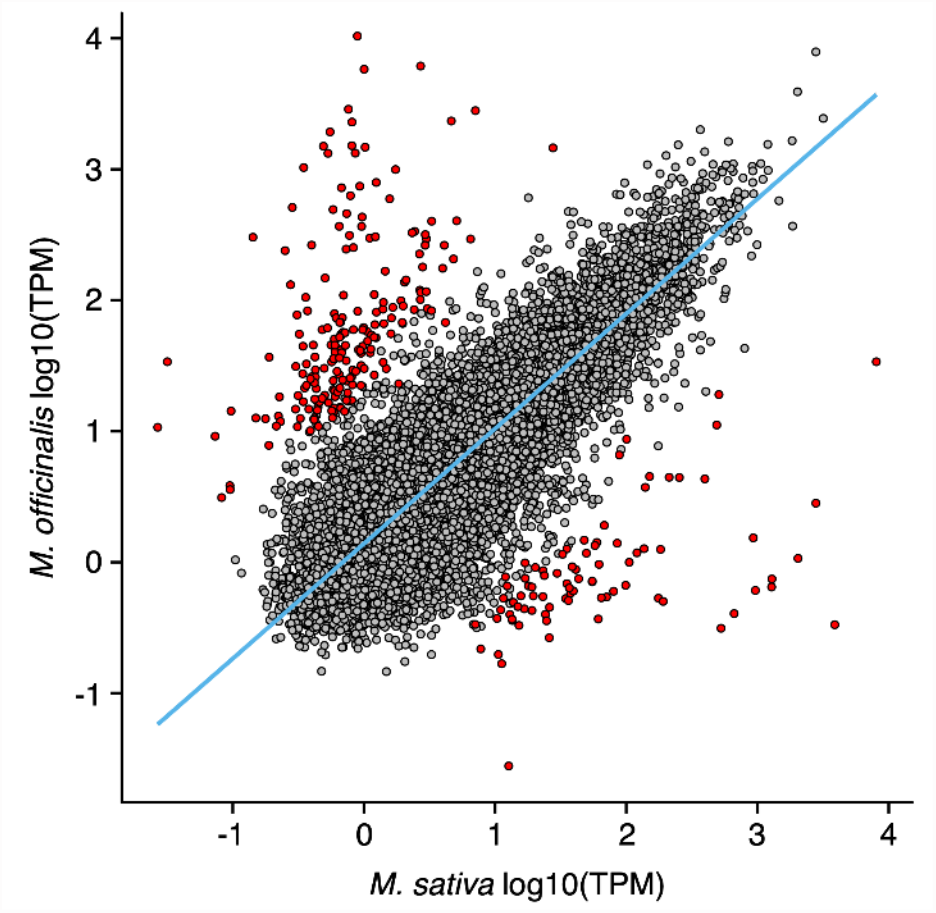
Correlation between transcript abundances of orthologous transcripts in *M. sativa* and *M. officinalis*. Each datapoint represents the transcript abundance of single-copy orthologous transcripts in *M. sativa* and *M. officinalis*. Red datapoints represent transcripts that are differentially abundant between the two species (|log_2_(fold change)| > 5, adjusted p-value < 0.01); all other datapoints are in grey. The blue line represents the robust linear regression line, calculated with the rlm function of the MASS package in R.

Many of the differentially-abundant conserved supertranscripts have annotated functions that suggest the encoded proteins may impact symbiotic nitrogen fixation. These include 21 supertranscripts annotated as encoding nodulins, which include 16 that are more abundant in *M. officinalis* and 5 that are more abundant in *M. sativa*. In addition, 32 supertranscripts encoding proteins predicted to be associated with transcription and translation activity were differentially abundant, with 24 more abundant in *M. sativa* and eight more abundant in *M. officinalis*. We also observed that several supertranscripts encoding proteins predicted to be involved in cell wall synthesis or modification were differentially abundant, with six more highly abundant in *M. sativa* and one more highly abundant in *M. officinalis*. Other differentially-abundant transcripts included those predicted to encode proteins involved in transport (17 transcripts), fatty acid biosynthesis (3 transcripts), flavonoid biosynthesis (3 transcripts), and aromatic compound biosynthesis (1 transcript). Given that this analysis compares two plant species with differing growth rates (**Table S2**), we cannot rule out that some of these transcriptomic differences may also reflect variances in nodule maturity and/or host metabolic activity at the time of harvest.

### NCR peptide diversity and expression profile

We previously observed that replacing the *bacA* allele of *S. meliloti* 2011 with the *bacA* alleles of the rhizobia *S. fredii* NGR234 or *R. leguminosarum* bv. viciae 3841 resulted in an inability to fix nitrogen with *M. sativa* while the ability to fix nitrogen with *M. alba* and *M. officinalis* remained ((24) and **Table S1**). We hypothesized that this was due to differences in the NCR peptide profiles of these species (24). To test this hypothesis, supertranscripts encoding NCR peptides were identified in the *M. sativa* and *M. officinalis* transcriptome assemblies using the SPADA pipeline (62). A total of 412 and 308 supertranscripts encoding NCR peptides were identified in the *M. sativa* and *M. officinalis* transcriptomes, respectively, accounting for ~0.5% of all supertranscripts in both assemblies (**Datasets S5 and S6**). The lower count of *NCR* transcripts in *M. officinalis* was offset by a higher median transcript abundance (58.6 transcripts per million [TPM] vs 99.1 TPM; p < 0.001; **Figure 5A**), resulting in *NCR* transcripts accounting for roughly 9% of the total nodule transcriptome in both species. In both *M. sativa* and *M. officinalis*, NCR peptides had median lengths of 38 residues, with approximately half of the NCR peptides containing between 30 and 40 residues (**Figure 5B**). Additionally, there was a roughly even number of four and six-cysteine NCR peptides expressed in both plant species, with the four-cysteine class of NCR peptides accounting for 51-55% of the *NCR* transcripts both in terms of number of NCR peptides and expression of *NCR* transcripts as measured by TPM. The NCR peptides from both hosts also showed broadly similar distributions of pI values between approximately 3 to 11, with one peak around a pI of 4 and another around pI 8 (**Figures 5C and 5D**). The pI pattern of the NCR peptides we observed is reminiscent of that reported for other legume species that induce an elongated branched morphology in their microsymbiont, including *M. sativa* and *M. truncatula* (86). Overall, at a global level, the property profiles of NCR peptides for *M. sativa* and *M. officinalis* were very similar, suggesting that the impact of different *bacA* alleles on symbiotic compatibility of *S. meliloti* with *M. sativa* is unlikely a consequence of global differences in the NCR peptide profiles of these plants and is more likely due to specific NCR peptides. Identifying which NCR peptides functionally correlate with symbiotic compatibility should be the focus of future studies.

**Figure 5.**
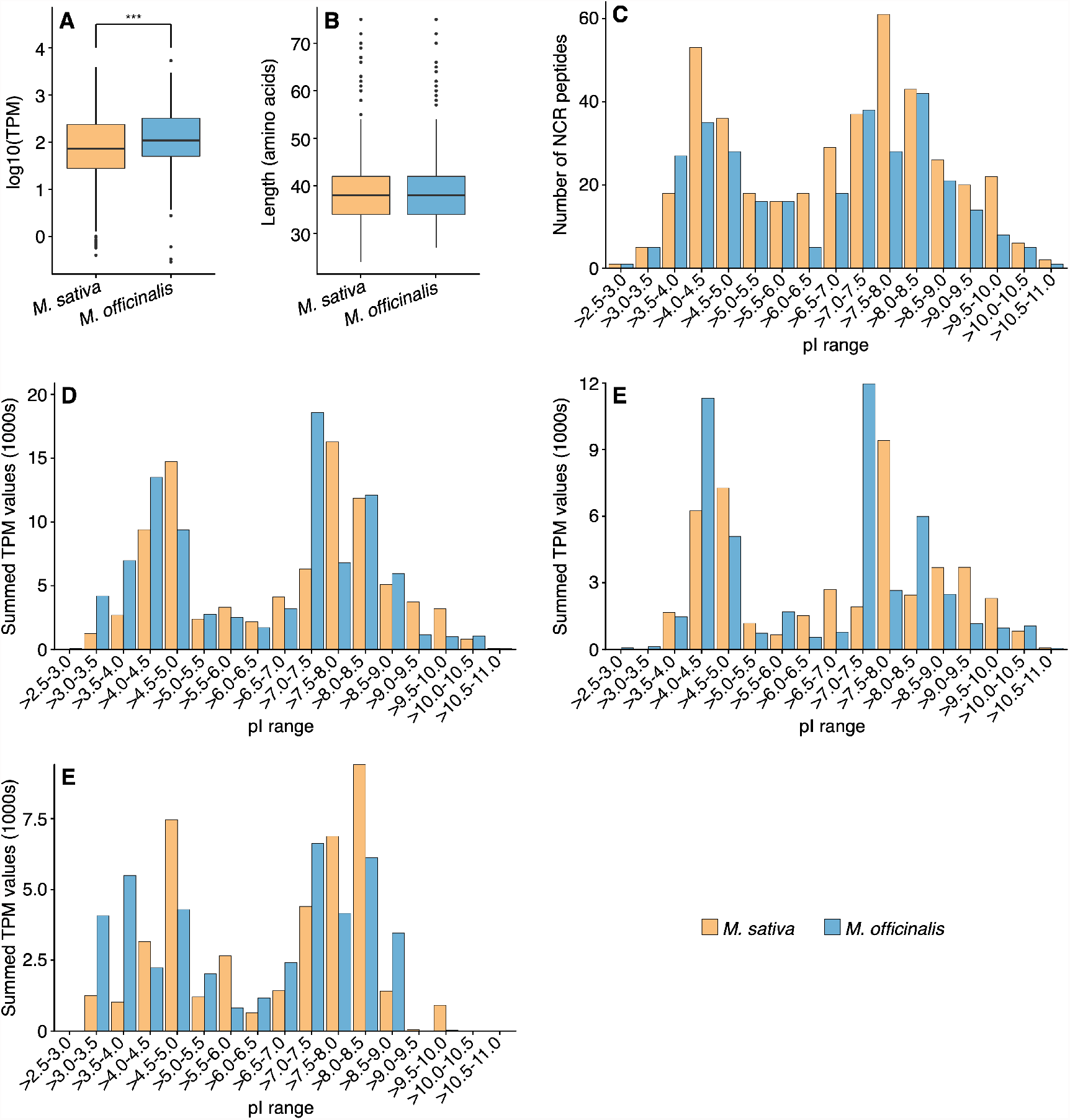
NCR peptide profiles of *Medicago sativa* and *Melilotus officinalis*. NCR peptides were predicted from the *M. sativa* (orange) and *M. officinalis* (blue) transcriptome assemblies, and the properties of the NCR peptides are shown in these graphs. (**A**) Box plots showing the distribution of the abundance (in transcripts per million, TPM) of *NCR* transcripts, based on triplicate samples. The difference in the distributions for the two species was statistically significant (p-value < 0.001; pairwise Wilcox test). (**B**) Box plots showing the distribution of the amino acid lengths of mature NCR peptides. No statistically significant difference in the distributions for the two species was detected. (**C**,**D**) Histograms showing the distributions of the isoelectric points (pI) for the mature NCR peptides. Histograms are based either on the number of NCR peptides with a given pI value (C) or the total abundance of the transcripts encoding NCR peptides with a given pI value (D). **(E**,**F)** Histograms showing distributions of pI for 4-cysteines (E) and 6-cysteines (F) mature NCR peptides based on total abundance of the transcripts encoding NCR peptides with a given pI value.

Despite the general similarity in the NCR peptide profiles of *M. sativa* and *M. officinalis*, a key difference emerges when examining the abundance of NCR peptides with extreme pI values; transcripts encoding highly cationic NCR peptides were more abundant in *M. sativa* while transcripts encoding highly anionic NCR peptides were more abundant in *M. officinalis* (**Figure 5D**). Previous work has shown that, in general, only cationic NCR peptides with a pI > 9.0 have antimicrobial activity (87), with anticandidal activity primarily limited to NCR peptides with a pI > 9.5 (88). Here, we observed that transcripts encoding highly cationic NCR peptides (pI > 9.0) were ~2.4-fold more abundant in *M. sativa* than *M. officinalis* (**Figure 5D**). Similarly, transcripts encoding NCR peptides with pI values > 9.5 were ~1.9-fold more abundant in *M. sativa* than *M. officinalis*. Notably, previous work indicated that 4.0% of *M. truncatula NCR* transcripts encode NCR peptides with pI values > 9.5, compared to only 1.8% in the *R. leguminosarum* bv. *viciae* symbiont *P. sativum* (86); this compares to 4.7% and 2.3% for *M. sativa* and *M. officinalis*, respectively (**Figure 5C**). Strikingly, when subdividing the NCR peptides with pI values > 9.5 into those with four or six cysteine residues, we observed that those with six-cysteines were ~27-fold more abundant in *M. sativa* than *M. officinalis* (**Figure 4E and 4F**). Considering these results, we hypothesize that the ability of the *R. leguminosarum bacA* allele to support symbiosis with *M. officinalis* and *P. sativum*, but not *M. sativa*, is a consequence of the elevated abundance of highly cationic (pI > 9.5) NCR peptides in *Medicago* nodules, with six-cysteine NCR peptides possibly being of particular significance. It may be that the BacA proteins of *S. fredii* and *R. leguminosarum* are less capable of transporting these NCR peptides, and consequently, strains with these BacA proteins may be more sensitive to the antimicrobial activities of these cationic NCR peptides.

## CONCLUSION

We report high quality nodule transcriptome assemblies for *M. sativa* cv. Algonquin and *M. officinalis* that we expect will serve as valuable resources for the legume research commumity. In particular, we expect that the availability of a nodule transcriptome for *M. officinalis* will help establish this plant as a secondary model system for studies of the symbiotic properties of *S. meliloti*.

We were particularly interested in using these transcriptomes to compare the properties of the NCR peptides encoded by both species. Despite predicting 33% more NCR peptides in *M. sativa* than *M. officinalis, NCR* transcripts accounted for roughly 9% of the transcriptome (based on TPM values) in both species. In general, the characteristics of the NCR peptides of *M. sativa* and *M. officinalis* were highly similar. However, transcripts encoding cationic NCR peptides with a pI > 9.5 were ~2-fold more abundant in *M. sativa* than in *M. officinalis*, and 27-fold more abundant when considering only six-cysteine NCR peptides. These results are consistent with previous observations that transcripts encoding cationic NCR peptides with a pI > 9.5 account for ~2-fold more *NCR* transcripts in *M. truncatula* compared to *P. sativum*. Cationic, but not neutral or anionic, NCR peptides display antimicrobial activity through disrupting the integrity of microbial membranes (89). It has been hypothesized that BacA provides protection against these NCR peptides by importing them into the cytoplasm and thus away from the membrane (90, 91). Considering that the BacA proteins of *S. fredii* and *R. leguminosarum* share less than 60% amino acid identity with the BacA protein of *S. meliloti*, it is reasonable to speculate that they have different substrate specificity and may be less capable of transporting cationic NCR peptides (24). If true, this could explain why the *bacA* alleles of *S. fredii* and *R. leguminosarum* can support symbiotic nitrogen fixation with *M. officinalis* but not *M. sativa*; the increased production of cationic NCR peptides in *M. sativa*, coupled with lower rates of import into the *S. meliloti* cytoplasm, could result in an accumulation of these peptides in the periplasm, resulting in a loss of viability and lack of nitrogen fixation (24). In future work, it will be interesting to test whether *S. meliloti* strains with different *bacA* alleles display differing sensitivities to these highly cationic NCR peptides, or differences in their abilities to transport these peptides.

## ACKNOWLEDGEMENTS

We thank Karen Ho and Neda Moradin from The Centre for Applied Genomics (Toronto, Canada) for helpful advice in planning the RNA-seq library preparation strategy. This research was enabled, in part, through computational resources provided by Compute Ontario (computeontario.ca) and Compute Canada (computecanada.ca). Funding for this research was provided by the Natural Sciences and Engineering Research Council of Canada (NSERC) through Discovery Grants to WAS and GCD.

