## Supplementary Tables and Figures for "Reference nodule transcriptomes for *Melilotus officinalis* and *Medicago sativa* cv. Algonquin"

**This PDF file includes:**

Tables S1 to S4
Figure S1
Legends for Datasets S1 to S6

**Other supplementary materials for this manuscript include the following:**

Datasets S1 to S6

**Table S1.** Shoot dry weights of *M. sativa* and *M. officinalis* plants inoculated with *S. meliloti* strains carrying various *bacA* constructs.

| *S. meliloti* strain | *bacA* allele | *M. sativa* (mg/plant) * | *M. officinalis* (mg/plant) * |
|---|---|---|---|
| Rm2011 | *S. meliloti* Rm2011 *bacA* | 52 ± 4 [a] | 115 ± 14 [a] |
| RmP3985 | Δ*bacA* | 6 ± 3 [b] | 5 ± 2 [c] |
| RmP3903 | *S. meliloti* Rm2011 *bacA* | 51 ± 8 [a] | 131 ± 13 [a] |
| RmP3905 | *S. fredii* NGR234 *bacA* | 8 ± 1 [b] | 49 ± 9 [b] |
| RmP3907 | *R. leguminosarum* bv. *viciae* 3841 *bacA* | 5 ± 3 [b] | 123 ± 21 [a] |
| Uninoculated | N/A | 4 ± 2 [b] | 4 ± 2 [c] |

* Data represent the mean ± the standard deviation of triplicate samples, with each sample consisting of between three and five plants. Different letters represent statistically unique groups ($\alpha < 0.05$) and were determined via one-way ANOVA followed by Tukey's HSD posthoc tests. Statistical analyses were performed separately for each plant species.

**Table S2.** Shoot dry weights of *M. sativa* and *M. officinalis* plants inoculated with wildtype *S. meliloti* Rm2011 and used for the transcriptome analyses.

| Plant | Number of Shoots | Dry Mass per Shoot (mg) |
|---|---|---|
| *M. sativa* replicate 1 * | 5 | 52 |
| *M. sativa* replicate 2 | 4 | 40 |
| *M. sativa* replicate 3 | 4 | 45 |
| *M. sativa* replicate 4 * | 4 | 42.5 |
| *M. sativa* replicate 5 | 4 | 92.5 |
| *M. sativa* replicate 6 * | 5 | 46 |
| *M. sativa* replicate 7 | 5 | 56 |
| *M. sativa* average | 4 | 53 |
| *M. officinalis* replicate 1 | 5 | 64 |
| *M. officinalis* replicate 2 * | 5 | 63.4 |
| *M. officinalis* replicate 3 * | 5 | 86 |
| *M. officinalis* replicate 4 | 4 | 90 |
| *M. officinalis* replicate 5 * | 5 | 76 |
| *M. officinalis* replicate 6 | 5 | 70 |
| *M. officinalis* replicate 7 | 5 | 88 |
| *M. officinalis* average | 5 | 77 |

* Asterisks denote replicates used for the transcriptome analyses.

**Table S3.** Summary statistics from the preprocessing of the Illumina reads.

| | *M. sativa* | *M. officinalis* |
|---|---|---|
| Raw paired-end reads | 202,126,321 | 138,919,600 |
| Paired-end reads corrected by Rcorrector * | 53,303,105 (26.3%) | 35,705,018 (25.7%) |
| Paired-end reads removed by Rcorrector * | 20,111,775 (10.0%) | 14,629,169 (10.5%) |
| Paired-end reads retained by Rcorrector * | 182,014,546 (90.0%) | 124,290,431 (89.5%) |
| Base pairs trimmed by Cutadapt † | 153,232,831 (0.4%) | 105,798,758 (0.3%) |
| Paired-end reads retained by Cutadapt † | 182,014,546 (100%) | 124,290,431 (100%) |
| Paired-end reads retained by Trimmomatic ¥ | 174,707,055 (96.0%) | 119,333,821 (96.0%) |

* Percentages are given relative to the number of raw paired-end reads.

† Percentages are given relative to the number of paired-end reads retained by Rcorrector or the number of base pairs presented in the paired-end reads retained by Rcorrector.

¥ Percentages are given relative to the number of paired-end reads retained by Cutadapt.

**Table S4.** Summary statistics from annotation of the compressed transcriptome assemblies.

| | *M. sativa* | *M. officinalis* |
|---|---|---|
| Total genes predicted by TransDecoder | 73,830 | 58,786 |
| Genes annotated via bi-directional blast with *Medicago truncatula* | 17,446 | 16,422 |
| Genes annotated using eggNOG-mapper | 15,091 | 10,303 |
| Genes annotated using the HMMs of the PFAM database | 843 | 1,493 |
| Genes annotated using the HMMs of the TIGRFAM database | 51 | 60 |
| Total number of annotated genes | 33,431 | 28,278 |
| Annotation rate (%) | 45 | 48 |

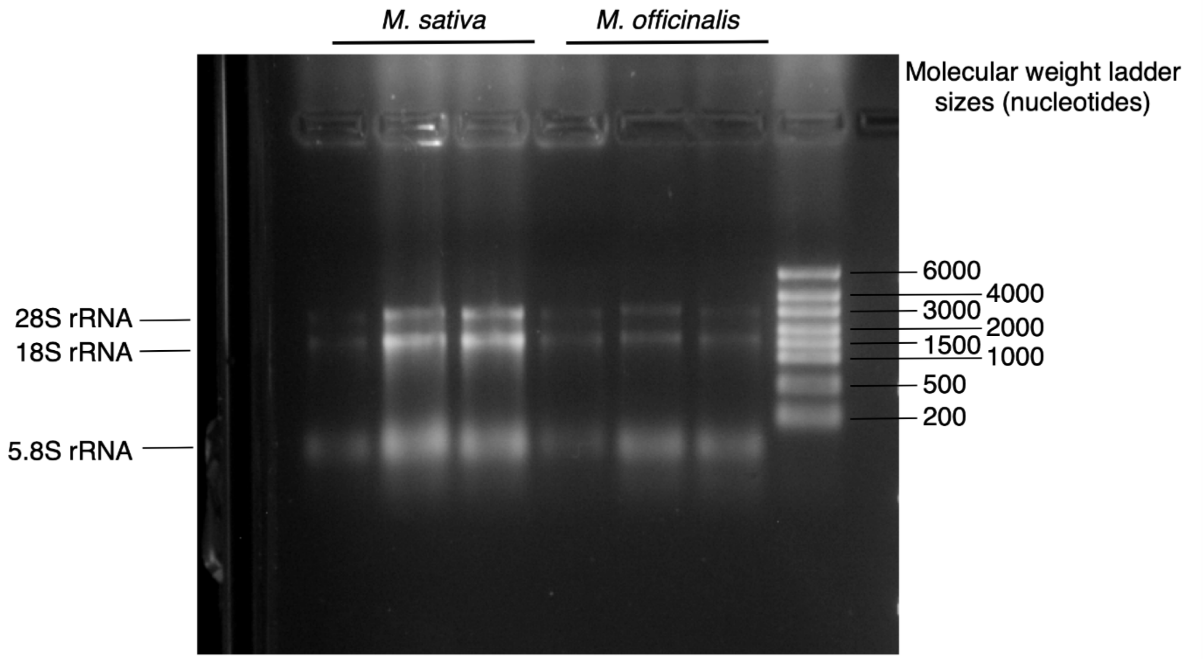


**Figure S1. Integrity of the RNA samples used for RNA-seq library preparation.** RNA samples were run on a MOPS-formaldehyde agarose gel and imaged. The bands corresponding to the 28S rRNA, 18S rRNA, and 5.8S rRNA are indicated. The lack of smearing indicates that the purified RNA was of high quality and not degraded.

### DATASETS

**Dataset S1.** Annotation of the *Medicato sativa* nodule transcriptome.

**Dataset S2.** Annotation of the *Melilotus officinalis* nodule transcriptome.

**Dataset S3.** Summary of the GO slim analysis of the *Medicago sativa* nodule transcriptome assembly.

**Dataset S4.** Summary of the GO slim analysis of the *Melilotus officinalis* nodule transcriptome assembly.

**Dataset S5.** NCR peptides predicted from the *Medicago sativa* nodule transcriptome assembly.

**Dataset S6.** NCR peptides predicted from the *Melilotus officinalis* nodule transcriptome assembly.
